# Fibroblast growth factor-21 induces skeletal muscle atrophy and increases plasma amino acids in female mice: a potential role for glucocorticoids

**DOI:** 10.1101/2023.06.27.546599

**Authors:** Karlton R. Larson, Devi Jayakrishnan, Karla A. Soto Sauza, Michael L. Goodson, Aki T. Chaffin, Arik Davidyan, Suraj Pathak, Yanbin Fang, Diego Gonzalez Magaña, Benjamin F. Miller, Karen K. Ryan

**Affiliations:** Department of Neurobiology, Physiology, and Behavior, College of Biological Sciences, University of California Davis, Davis, California 95616; Department of Biological Sciences, California State University Sacramento,6000 J Street, Sacramento, CA, 95819, USA; Aging & Metabolism Program, Oklahoma Medical Research Foundation, Oklahoma City, Oklahoma, USA; Oklahoma City Veterans Affairs Medical Center, Oklahoma City, Oklahoma, USA

**Keywords:** FGF21, stress, HPA axis, glucocorticoid, muscle, protein restriction

## Abstract

**Background:** Fibroblast growth factor-21 (FGF21) is an intercellular signaling molecule secreted by metabolic organs, including skeletal muscle, in response to intracellular stress. FGF21 crosses the blood brain barrier and acts via the nervous system to coordinate aspects of the adaptive starvation response, including increased lipolysis, gluconeogenesis, hepatic fatty acid oxidation, and activation of the hypothalamic-pituitary-adrenocortical (HPA) axis. Given its beneficial effects for hepatic lipid metabolism, pharmaceutical FGF21 analogues are in clinical trials treatment of fatty liver disease. We predicted pharmacologic treatment with FGF21 increases HPA axis activity and skeletal muscle glucocorticoid signaling and induces skeletal muscle atrophy in mice.

**Methods:** We treated male and female mice with FGF21 or saline, delivered either peripherally or directly to the brain, to determine its effect on skeletal muscle. To identify metabolic pathways affected by FGF21, we analyzed untargeted primary metabolites measured in plasma by GCTOF-MS. To determine mechanisms underlying sex-and FGF21-dependent changes in muscle mass, we measured hormonal and molecular mediators of muscle protein synthesis and degradation. We performed stable isotope labeling with deuterium oxide to directly measure muscle protein synthesis.

**Results:** A short course of systemic FGF21 treatment decreased muscle protein synthesis (*P* < 0.001) and reduced tibialis anterior weight (*P* < 0.05); this was driven primarily by its effect in female mice (*P* < 0.05). Similarly, intracerebroventricular FGF21 reduced TA muscle fiber cross sectional area (*P* < 0.01); this was more apparent among female mice compared to male littermates (*P* < 0.05). In agreement with the reduced muscle mass, the topmost enriched metabolic pathways in FGF21-treated females were related to amino acid metabolism, and the relative abundance of plasma proteinogenic amino acids were increased up to three-fold (*P* < 0.05). FGF21 treatment increased hypothalamic *Crh* mRNA (*P* < 0.01), plasma corticosterone (*P* < 0.01), and adrenal weight (*P* < 0.05), and increased expression of glucocorticoid receptor target genes known to reduce muscle protein synthesis and/or promote degradation including *Foxo1*, *Redd1*, and *Klf15* (P < 0.05). Again, these changes were driven primarily by effects of FGF21 in females (*P* < 0.05).

**Conclusions:** FGF21 increased plasma amino acids and decreased skeletal muscle mass, together with activation of the HPA axis and glucocorticoid receptor target genes in skeletal muscle—and female mice were more sensitive to all these outcomes. Given the proposed use of FGF21 analogues for the treatment of metabolic disease, the study is both physiologically relevant and may have important clinical implications.

## Introduction

An adequate supply of amino acids is needed for essential physiological processes, including protein synthesis, hormone and neurotransmitter production, and fuel metabolism (1). In the face of dietary protein or caloric restriction, and even during starvation, circulating concentrations of amino acids remain remarkably stable (2,3). The ability to maintain circulating amino acid concentrations, across a range of environmental and internal perturbations, highlights the presence of physiologic mechanisms that control systemic amino acid homeostasis. We and others find that changes in feeding behavior and macronutrient selection represent one such mechanism (4–10). And when behavioral modifications are unavailable or insufficient, structural and functional proteins of the body can be accessed to provide amino acids in times of need (11).

Skeletal muscle serves as the largest accessible reservoir of protein and amino acids in the body. In response to starvation or other metabolic stressors, the skeletal muscle protein pool can be accessed to provide amino acids for use as fuel or for cellular maintenance and repair (12). To accommodate these demands, the dynamic balance between protein synthesis and protein degradation is modified to favor muscle atrophy; this metabolic adaptation to starvation is largely accomplished by actions of the hypothalamic-pituitary-adrenal (HPA) axis (12,13). Glucocorticoid receptor (GR) signaling in skeletal muscle upregulates the expression of key transcription factors leading to muscle catabolism, including krupple-like factor-15 (*Klf15*) and the forkhead box proteins (*Foxo*’s) (14–18). Additionally, GR activity represses protein synthesis by increasing gene expression for Regulated in development and DNA damage response-1 (*Redd1*). Both Redd1 and Klf15 inhibit the master regulator of protein synthesis, mechanistic target of rapamycin complex-1 (mTORC1) (16,19). The cumulative effect of GR activation is increased free amino acid availability in skeletal muscle and plasma. Chronic exposure to excess glucocorticoids thereby leads to skeletal muscle atrophy (13,20–22).

Fibroblast growth factor-21 (FGF21) is an intercellular signaling molecule that is produced by various organs and tissues in response to intracellular stress (23). It was initially identified as a hepatokine, secreted into circulation by the liver during a prolonged fast (23–25). Newer data revealed that protein or amino acid deficit, rather than fasting or starvation *per se*, drives FGF21 secretion (26). FGF21 signals via the FGF-receptor 1 (FGFR1), a receptor tyrosine kinase, and its obligatory co-receptor β-Klotho (KLB) (27). FGFR1 has low expression in liver (28). Accordingly, FGF21 is thought to act in an endocrine manner to coordinate several aspects of the adaptive starvation response (29,30), including lipolysis, hepatic fatty acid oxidation (31) and gluconeogenesis (32), together with activation of the HPA axis (32,33). Collectively, these data support the possibility that FGF21 also increases systemic amino acid availability by accessing skeletal muscle protein and amino acids via the HPA axis.

FGF21 is a promising therapeutic target for the treatment of hepatic steatosis, because it potently reduces liver triglycerides in mice, monkeys, and humans (23,25,34–37). Recent work by ourselves and others revealed that the benefit of FGF21 for hepatic lipid metabolism is sex dependent. FGF21 decreased liver triglycerides in obese male mice, but this was abrogated among obese female littermates (38,39). Thus, sex is an important biological variable influencing FGF21 physiology. Similarly, it is well established that sex is an important variable influencing stress physiology. Both basal HPA axis tone and the glucocorticoid response to stressors is increased in female rodents and women compared to male counterparts (40–42). In addition, female skeletal muscle is more sensitive to glucocorticoid-induced atrophy (12,43).

Here we tested the hypothesis that pharmacologic administration of FGF21 activates the HPA axis, resulting in increased plasma amino acids and skeletal muscle atrophy. Because aspects of FGF21 and HPA axis biology are sex dependent, we included both male and female mice in our experimental design. The study is both physiologically relevant and may have important implications for the proposed clinical use of FGF21 analogues in the treatment of metabolic disease.

## Methods

### Animals

All animal experiments were approved by the Institutional Care and Use Committees of the University of California, Davis. Age-matched male and female C57BL/6J mice (n = 3-13/group) were obtained from The Jackson Laboratory or were bred in-house, for up to two generations removed from founders obtained from The Jackson Laboratory. Because therapeutic interest in pharmacologic FGF21 stems from its beneficial effects for obesity-associated metabolic disease, all experiments used diet-induced obese (DIO) mice.

### FGF21 administration

For systemic administration, mice received 0.2mg/kg/day FGF21 or an equivalent volume of saline. Twice daily intraperitoneal (i.p.) injections (0.1mg/kg/injection) were performed for 12 days on mice included in Figure 1. Once daily subcutaneous (s.c.) injections (0.2mg/kg/injection) were performed for 1, 4, 8, 15, and 32 days for mice included in Figure 6. Mice were single-housed on a 12-hour light, 12-hour dark cycle in a temperature (20-22° C) and humidity-controlled vivarium with *ad libitum* access to food and water.

**Figure 1.**
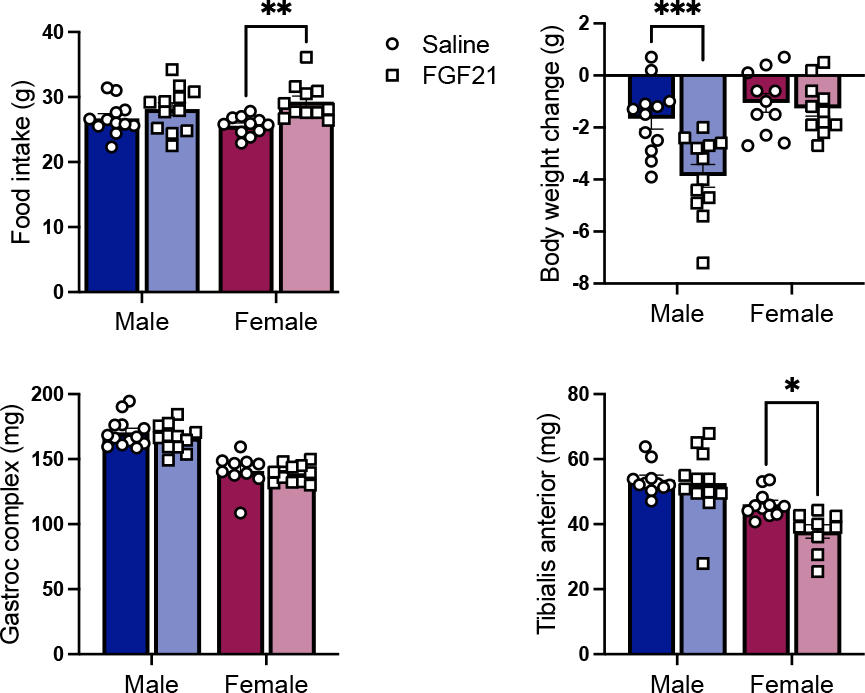
*FGF21-induced muscle atrophy in female mice.* FGF21-treated mice (0.2 mg/kg/d i.p. for 12 days) ate more than saline-treated controls [**A**, P (treatment) < 0.01]. FGF21 caused body weight loss, in a sex-dependent manner [**B**, P (treatment x sex) < 0.01]. FGF21 had no effect on gastrocnemius-soleus complex weight [**C**], but it significantly decreased wet weight of the tibialis anterior [**D**, P (treatment) < 0.01]. Tukey’s HSD posthoc test, **\***<0.05, **\*\***<0.01; **\*\*\***< 0.001, N= 9-13/ group.

For intracerebroventricular (i.c.v.) administration (Figure 4), mice received continuous infusion of FGF21 (0.4 μg/d) or an equivalent volume of saline for 13 days. We chose this dose because it is effective to activate brain FGF-receptor signaling in rodents, without efflux to the periphery (32,44). First, anesthetized mice were fitted with an i.c.v. cannula (Brain Infusion Kit 3, Alzet Corporation, Palo Alto, CA) fitted to an Alzet osmotic minipump (0.25 μl/h, model 1002), which administered either FGF21 or saline continuously for 13 days. Brain infusion cannulas were placed using a stereotaxic apparatus (Kopf Instruments, Kujunga, CA) with tips in the lateral cerebral ventricle using the following coordinates: 0.7 mm posterior to bregma, 1.2 mm lateral to the midsagittal suture, and to a depth of 2.5 mm from the brain surface. The connected minipumps were inserted in a subcutaneous pocket on the mouse’s flank, and the cannulas were secured to the skull and wound closed using vetbond (3M. St. Paul, MN). The cannulas were secured by glue. The animals received 5 mg/kg meloxicam after surgery and on the first two postoperative days. The location of the cannula was confirmed at the end of the experiment by methylene blue staining.

### Diets

High fat diet (HFD, D12492) was manufactured by Research Diets (New Brunswick, NJ) and based on the American Institute of Nutrition Rodent Diets growth formula, AIN-93G(45) containing 20% kcal from protein, 20% kcal from carbohydrate, and 60% kcal from fat.

### Tissue Collection

Mice were euthanized with a pentobarbital injection. Whole trunk blood was collected in chilled EDTA-coated tubes either following decapitation or using cardiac puncture and centrifuged at 3000 x g for 15 minutes at 4° C. Plasma was aliquoted and stored for later use at −80° C. The tibialis anterior (TA) muscle, gastrocnemius-soleus complex, and inguinal white adipose tissues (iWAT) were dissected free of connective tissue, weighed, and frozen in dry ice-cooled isopentane. Muscles from the right limb were pinned to cork board prior to freezing for histological use.

### Muscle fiber cross sectional area (CSA)

We cut serial cross sections (10 μm) from the TA and soleus using a Leica CM 3050S cryostat. To determine cross-sectional area (CSA), we fixed TA muscle sections in cold acetone for 5 min at −20°C, followed by three 5-min washes with phosphate-buffered saline with 0.1% Tween 20. We then incubated the sections with mouse-on-mouse block (MKB-2213 VECTOR laboratories) in phosphate-buffered saline (60 μl: 2.5 mL) for one hour at RT. We blocked sections in 5% normal goat serum in phosphate-buffered saline with 0.1% Tween 20 (blocking buffer) for 30 min at room temperature (RT) and then incubated in primary antibody for laminin protein (1:500, Sigma catalog no. L93931) overnight at 4°C. After incubation in primary antibody, we incubated sections in goat-anti-rabbit AlexaFluor 647 secondary antibody for 30 min at RT, and then cover slipped using ProLong Gold Antifade reagent (Life Technologies, catalog no. P36930). We imaged the slides using Leica DFC300 light microscope at 2x objective and analyzed using Cell-Pose in ImageJ. Fibers from four regions of a single section were analyzed per muscle, per animal.

### Stable Isotope Labeling & Protein Synthesis

For the data shown in Figure 6, newly synthesized proteins were labeled using deuterium oxide (D2O) according to previously published guidelines (46,47). Briefly, prior to starting chronic FGF21 injections, mice were first given an i.p. injection of 99% D2O equivalent to 5% of the body water pool. Mice were subsequently provided *ad libitum* access to 8% D2O enriched drinking water for the duration of the experiment. Male and female mice were sacrificed at 1, 4, 8, 15, and 32 days after initial bolus of D2O (n=3/ sex/ treatment/ time point) (48).

At euthanasia, muscles were weighed and rapidly frozen in liquid nitrogen. For analysis of tracer enrichment, TA was powdered using a liquid nitrogen-cooled mortar and pestle. Tissue was homogenized and fractionated to obtain the myofibrillar protein fraction and plasma was prepared as in (46,49) to determine the precursor pool enrichment. Derivatized alanine was analyzed on an Agilent 7890A Gas Chromatography coupled to an Agilent 5975 Mass Spectrometry as previously described (49–51). The deuterium enrichments of both the protein (product) and the precursor were used to calculate fraction new: Fraction new = Eproduct /Eprecursor, where the Eproduct is the enrichment (E) of protein-bound alanine and Eprecursor is the calculated maximum alanine enrichment from equilibration of the body water pool. The fraction new data were then plotted across the time points to calculate k, (1/day) using a one-phase association (48).

### FGF21

Endotoxin-free, recombinant human FGF21 (hFGF21; ProSpecBio, Rehovot, Israel) was first dissolved in H2O, according to the manufacturer’s instructions, and was further diluted in 0.9% sterile saline.

### Gene Expression

Tissue was collected from the mice at euthanasia, after 12 days of i.p. FGF21 or saline treatment, and frozen in isopentane for later analysis. The muscles were first powdered using a mortar and pestle chilled with liquid nitrogen and homogenized in QIAzol Lysis Buffer using a Bead Beater. Total RNA was isolated from the homogenate using the RNeasy mini kit. Complementary DNA was synthesized using the high-capacity complementary DNA reverse transcription kit (Life Technologies/ThermoFisher, Carlsbad, CA). Gene expression analysis was performed in duplicate with TaqMan gene expression assays and utilizing a Roche LightCycler 480. Samples with poor technical replicates (>0.5 Ct difference) were excluded from further analysis. Expression of DNA-damage inducible transcript 4 (aka REDD1) (*Ddit4*; Mm00512504_g1), corticotropin releasing hormone (*Crh*; Mm01293920_s1), beta-klotho (*Klb*; Mm00473122_m1), fibroblast growth factor receptor 1c (*Fgfr1*; Mm00438930_m1), forkhead box 01 (*Foxo1*; Mm00490671_m1), muscle RING-finger protein-1 (*Trim63*; Mm01185221_m1), muscle atrophy F-box (*Fbxo32*; Mm00499523_m1), and Krüppel-like factor-15 (*Klf15*; Mm00517792_m1), was normalized to the housekeeping gene 18S rRNA (*Rn18s; Mm04277571_*s1) using the 2 ^τιτιCt^ method.

### Plasma hormones

Circulating corticosterone and insulin-like growth factor-1 (IGF-1) was measured by the University of California, Davis Mouse Metabolic Phenotyping Center (MMPC). Corticosterone was measured using radioimmunoassay (RIA, MP Biomedicals, Orangeburg, NY) and IGF-1 was measured using an enzyme-linked immunosorbent assay (R & D Systems, Minneapolis, MN). Samples flagged by the MMPC for poor quality were removed from further analysis.

### Plasma metabolomics

Plasma was submitted to the West Coast Metabolomics Center at the University of California, Davis for untargeted primary metabolite analysis by automated liner exchange cold injection system (ALEX-CIS) gas chromatography time of flight (GCTOF) mass spectroscopy (MS). Data were acquired as previously described (38,52). Metabolomics data were analyzed using MetaboAnalyst 5.0 and the MetaboAnalystR package (version 3.3.0) in R (version 4.3.0). Sample concentrations in plasma collected from saline or FGF21 treated mice, separated by sex, were normalized using sum normalization and data were scaled using mean centering before using the provided KEGG compound identifiers to perform KEGG pathway Quantitative Enrichment Analysis. Metabolomics data are available at the NIH Common Fund’s National Metabo-lomics Workbench data repository, https://www.metabolomicsworkbench.org, under assigned Study ID STXXXXX (53). The data can be accessed directly via their Project DOI: XXXX. This work is supported by NIH grant U2C-DK119886.

### Statistical analysis

The remaining data were analyzed using GraphPad Prism (GraphPad Software, La Jolla, CA) and/or SigmaStat (Systat Softward, San Jose, CA) by T-test or 2-way ANOVA. Planned comparisons were made using the *Tukey* honestly significant difference test. Data are plotted as means +/- standard error of the mean unless otherwise noted.

## Results

### FGF21 and energy balance

An extensive literature reports that FGF21 reduces body weight and body fat in male mice via increased energy expenditure, and this is often associated with increased food intake (29,34,44,54). In agreement with this literature, mice treated with FGF21 (0.2mg/kg/day, i.p.) consumed more calories than saline treated controls, over the 12 days of treatment [p (treatment) < 0.01] **(Fig. 1A).** The effect of FGF21 on body weight and body fat loss depends on sex, such that FGF21-induced body weight and fat loss is more apparent in male mice compared to female littermates (38). Likewise in this study, FGF21-treated mice lost weight compared to saline-treated controls, in a sex-dependent manner [p (treatment x sex) < 0.01]. FGF21 induced weight loss was more apparent in males (Tukey, p< 0.001) **(Fig. 1B)**.

### FGF21 and skeletal muscle atrophy

To determine the effect of chronic i.p. FGF21 treatment on skeletal muscle mass, mice were euthanized, and we collected and weighed the gastrocnemius-soleus complex and TA muscles. Muscle mass was smaller in female mice compared to male littermates, as expected given their smaller body size [P (sex) < 0.001] **(Fig. 1C, D)**. There was no effect of FGF21 treatment on gastrocnemius-soleus complex weight **(Fig. 1C)**. However, TA weight was significantly less in FGF21-treated groups [P (treatment) < 0.05], and the statistical significance was primarily driven by the effect of FGF21 in female mice [Tukey, P < 0.05] **(Fig. 1D)**.

### FGF21 and plasma metabolites

To identify metabolic pathways affected by FGF21, we analyzed untargeted ‘primary metabolites’ (carbohydrates and sugar phosphates, amino acids, hydroxyl acids, free fatty acids, purines, pyrimidines, aromatics, exposome-derived chemicals) in plasma from DIO mice treated for 12 days with FGF21 (0.2mg/kg/d, i.p.) or saline. In agreement with the effect of FGF21 to induce muscle atrophy in female mice, KEGG pathway Quantitative Enrichment Analysis revealed four of the top-five most enriched pathways in females were related to amino acid metabolism **(Fig. 2B).** These included: ‘Folate metabolism’, ‘Catecholamine biosynthesis’, ‘Thyroid hormone synthesis’, ‘Tyrosine metabolism’, and ‘Arachidonic acid metabolism’. In agreement with our recent report that the effect of FGF21 to reduce hepatic and adipose tissue triglycerides is more apparent in male mice compared to female littermates (38), the top-five most enriched pathways in males were related to lipid metabolism **(Fig. 2A).** These included: ‘Steroidogenesis’, ‘Steroid biosynthesis’, ‘Bile acid biosynthesis’, ‘Glycerolipid metabolism’, and ‘Alpha Linolenic Acid and Linoleic acid metabolism’. Lastly, FGF21-treated female mice had greater relative concentration of the proteinogenic amino acids in plasma compared to saline-treated controls [P (treatment) < 0.05] **(Fig. 2D).** Thus, pharmacologic treatment with FGF21 alters amino acid metabolism and increases plasma amino acid availability in female mice.

**Figure 2.**
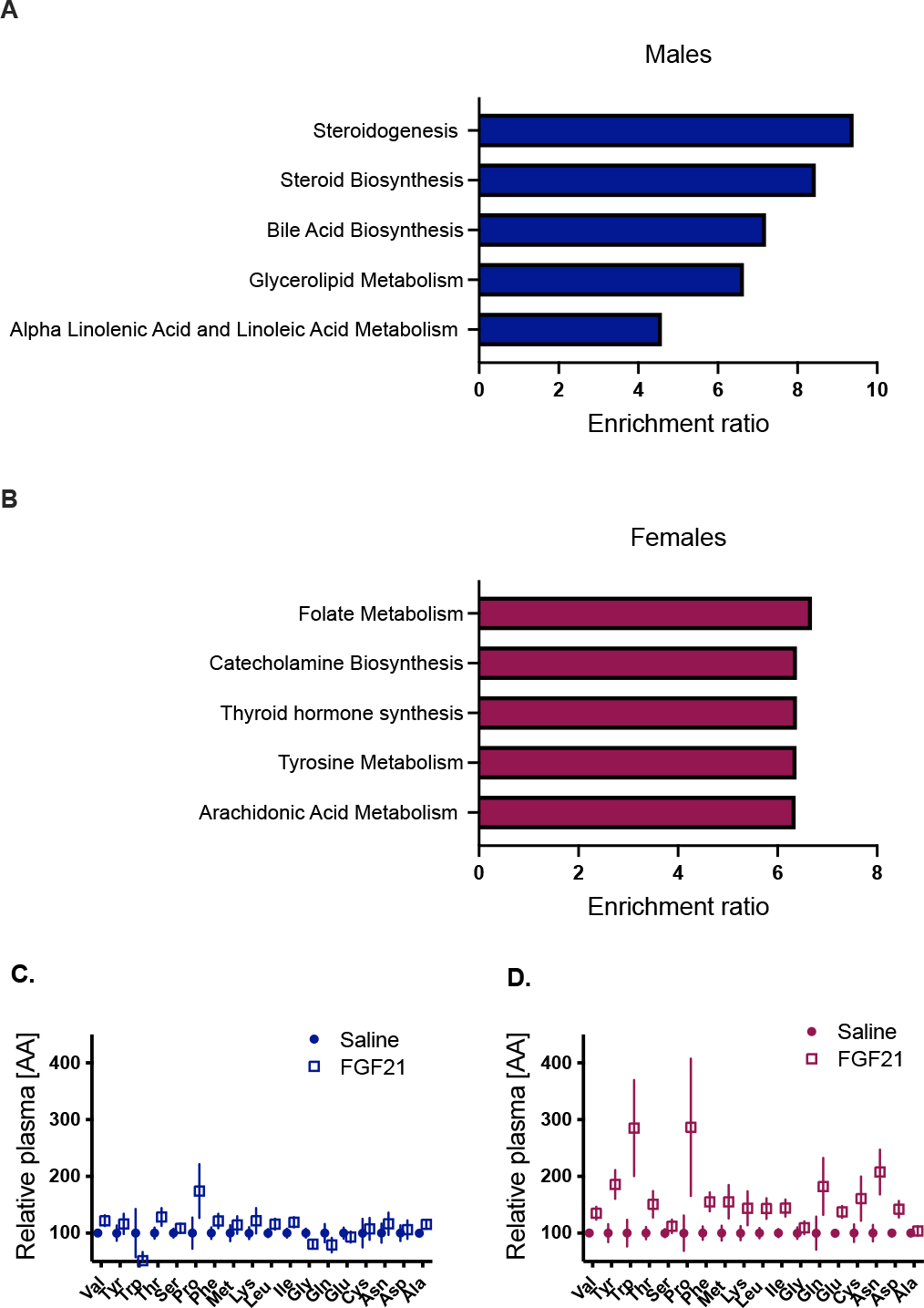
*FGF21 and the plasma metabolome.* The top 5 most enriched KEGG pathways from FGF21 (vs saline) treated males were related to lipid metabolism (**A**). 4 of the top 5 most enriched KEGG pathways from FGF21 (vs saline) treated females were related to amino acid metabolism (**B**). FGF21 increased the relative abundance of proteinogenic amino acids in plasma, and this was more apparent in females [**D**, P (treatment) < 0.05] compared to male littermates [**C**]. N= 9-13/ group.

### FGF21 treatment and the HPA axis

Because chronic glucocorticoid exposure both induces muscle atrophy and increases circulating amino acids, we measured indices of HPA axis activity. First, we dissected hypothalamus from mice treated for 12 days with FGF21 (0.2mg/kg/day, i.p.) or saline. To confirm hypothalamic expression of the FGF21 receptor complex, we measured mRNA for *Fgfr1* and *Klb.* Expression of these genes was not affected by sex or by FGF21 treatment **(Fig. 3A** and **3B)**. FGF21-treated mice showed greater expression of hypothalamic corticotropin releasing hormone (*Crh*) mRNA [P (treatment) < 0.05], and the statistical significance was driven primarily by the effect in female mice [Tukey *post hoc* P < 0.05] **(Fig. 3C)**. Next, we measured corticosterone in plasma, by RIA. As expected, female mice had higher plasma corticosterone compared to male littermates [P (treatment) < 0.001]. FGF21-treated mice showed higher circulating corticosterone compared to controls [P (treatment) < 0.01] **(Fig. 3D)**, and this effect was more apparent in female mice [Tukey *post hoc* P < 0.01]. Corticosterone suppresses plasma insulin-like growth factor-1 (IGF1), which we also observed [P (treatment) < 0.001] **(Fig. 3E)**. Again, the statistical significance was driven primarily from the effect in female mice [Tukey *post hoc*, p< 0.001]. As an index of chronic HPA axis activity (33,55–57), we dissected, cleaned, and weighed the adrenal glands. As expected, female mice had larger adrenal glands than male littermates [P (sex) < 0.001]. Also, FGF21-treated mice had larger adrenal glands compared to saline-treated controls [P (treatment) < 0.05] (**Fig. 3F**) (58). These findings are consistent with increased HPA axis activity in FGF21-treated mice, which is more apparent among females.

**Figure 3.**
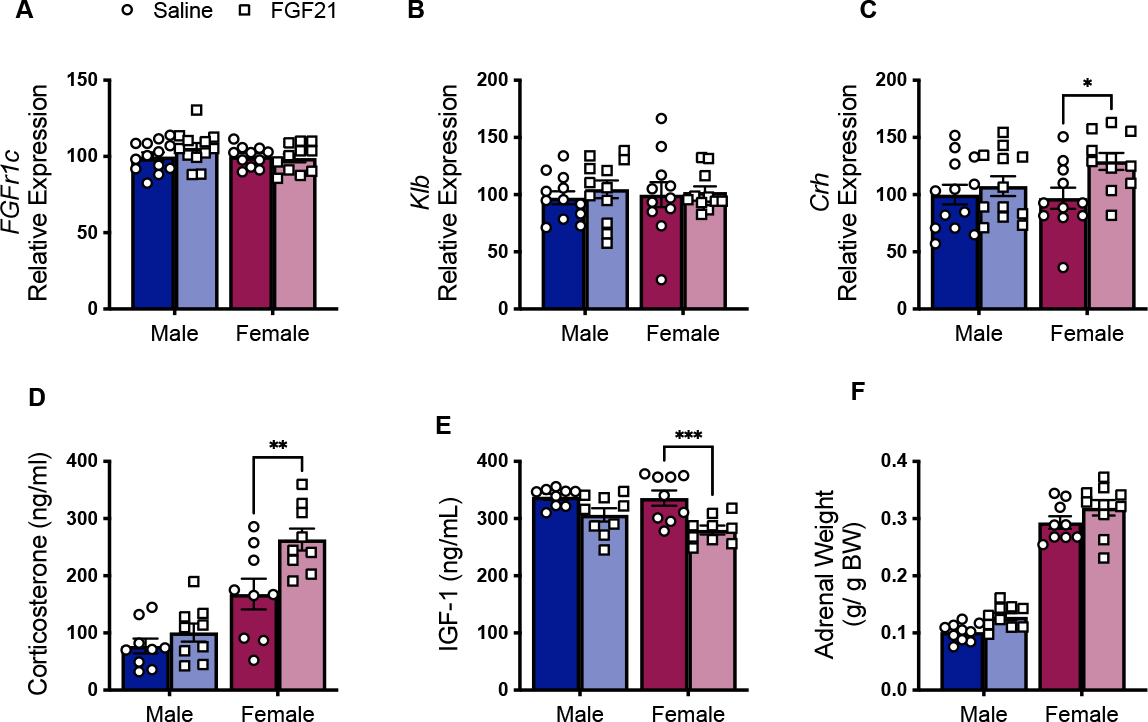
*FGF21 increased HPA axis activity.* FGF21-receptor complex mRNAs (*Fgfr1c* and *Klb*) were expressed in male and female hypothalamus and were not affected by FGF21 treatment (0.2mg/kg/d i.p. for 12 days) (**A, B**). FGF21-treated mice had increased expression of corticotropin releasing hormone (*Crh*), and increased plasma corticosterone [**C**-**D**, P (treatment) < 0.05]. FGF21 reduced plasma insulin-like growth factor 1 (IGF-1) [**E**, P (treatment) < 0.001]. FGF21-treated mice had larger adrenal glands [**F**, P (treatment)< 0.001]. Tukey’s HSD posthoc test, **\***<0.05, **\*\***<0.01; **\*\*\***< 0.001, N= 9-13/ group.

### Intracerebroventricular administration of FGF21 and muscle atrophy

FGF21 acts directly in the hypothalamus to activate the HPA axis and increase plasma glucocorticoids (32,33). To isolate the cell non-autonomous effects of FGF21 on skeletal muscle, we delivered FGF21 or its saline vehicle directly to the brain of DIO male and female littermates via osmotic pump to the lateral ventricle (0.4 μg/ day), for 13 days. As before, FGF21-treated mice lost more weight than saline-treated controls [P (treatment) < 0.01] (**Fig 4A**) and the statistical significance was driven primarily by the effect in males [Tukey, P < 0.05]. FGF21-treated male mice had less fat in the inguinal depot compared to saline-treated controls [t-test, P < 0.05] (**Fig 4B**). As expected, female mice had smaller TA muscles than male mice [P (sex) < 0.001] (**Fig 4C**). Lastly, i.c.v. FGF21-treated mice had smaller muscle fiber cross sectional area compared to saline-treated controls [P (treatment) < 0.01] (**Fig 4D**) which was more apparent in female mice [Tukey, P< 0.05].

**Figure 4.**
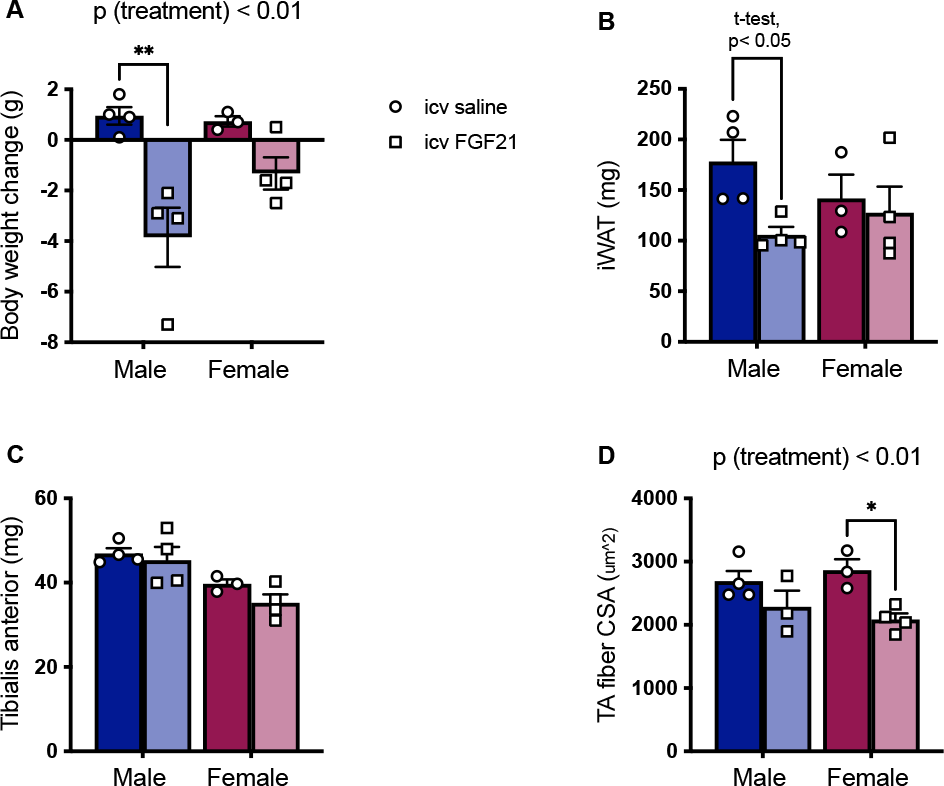
*Intracerebroventricular FGF21 reduces muscle fiber cross sectional area in female mice.* FGF21-treated mice (0.4 ug/d i.c.v. for 13 days) lost more weight than saline-treated littermates [**A**, P (treatment) < 0.01]. FGF21-treated male mice had smaller inguinal white adipose tissue (iWAT) depots than saline-treated controls [**B,** t-test]. Female mice had smaller tibialis anterior (TA) muscles than males [**C,** P (sex) < 0.001]. FGF21 reduced muscle fiber cross sectional area (CSA) [**D**, P (treatment) < 0.01]. Tukey’s HSD posthoc test, **\***<0.05, **\*\***<0.01, N= 3-4/ group.

### FGF21 treatment increases transcription of glucocorticoid target genes in TA muscle

Glucocorticoid receptors (GR) acts as a transcription factor for gene networks that induce skeletal muscle atrophy. GR activity increases the transcription of genes that reduce muscle protein synthesis and promote degradation (12,15,22). We measured mRNA for GR target genes in TA collected from the mice in Figure 1. FGF21-treated mice had greater expression of mRNA for DNA-damage-inducible transcript-4 (aka *Ddit4* or *Redd1*) [P (treatment) < 0.05] and Krüppel-like factor 15 (*Klf15*) [P (treatment) < 0.05] **(Fig. 5A** and **5B),** which reduces muscle protein synthesis by inhibiting mTORC1 (59). FGF21-treated mice had greater relative expression of mRNA for Forkhead-box O1 (*Foxo1*), and this was sex-dependent [P (treatment x sex) < 0.01] **(Fig. 5C)**. All these outcomes were more apparent among female mice [Tukey post hoc P < 0.05]. In turn, FoxO1 and KLF15 regulate the transcription of the E3 ligases muscle RING-finger protein-1 (MuRF1) and muscle atrophy F-box (MAFbx), which are a part of the ubiquitin-proteasome system, the primary intracellular degradation pathway (12). However, there was no significant effect of FGF21 treatment on either of these downstream targets **(Fig. 5D** and **5E).** There was no effect of FGF21 on skeletal muscle GR mRNA **(Fig. 5F)**. Collectively, these data support a mechanistic role for skeletal muscle GR-signaling and decreased protein synthesis in FGF21-induced skeletal muscle atrophy in female mice.

**Figure 5.**
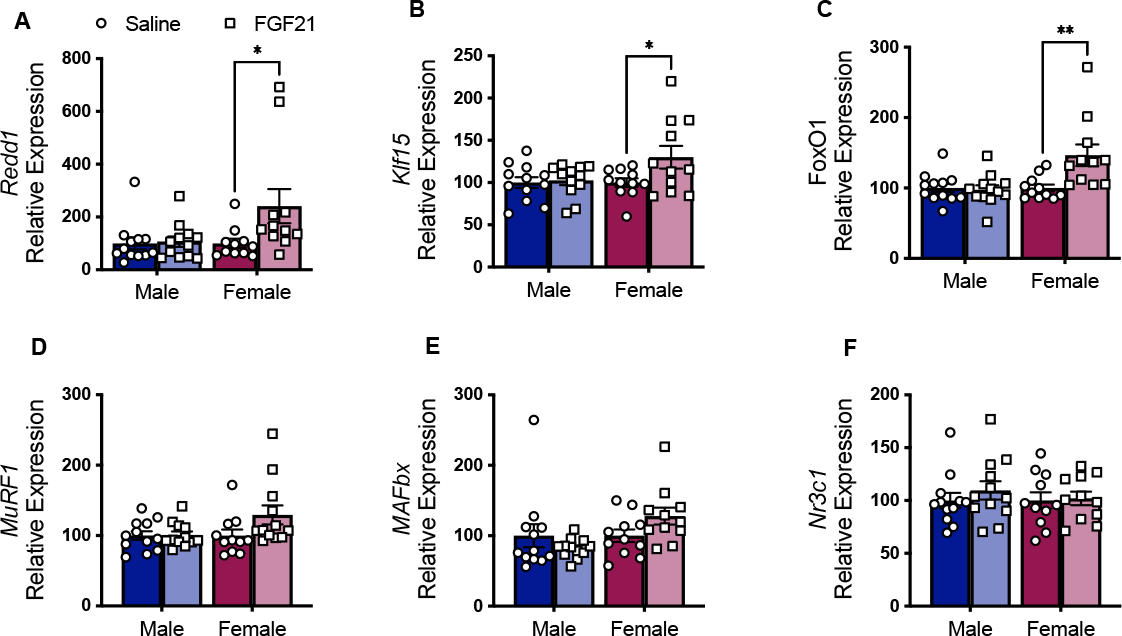
*FGF21 increased the expression of GR target genes in skeletal muscle of female mice.* i.p. FGF21 (0.2 mg/kg/d for 12 days) increased the expression of GR target genes *Redd1* and *Klf15* [**A-B** P (treatment)< 0.05]. FGF21 increased the relative expression of GR target gene *Foxo1* in a sex-dependent manner [**C** P (treatment x sex) < 0.01]. There was no significant effect of FGF21 on expression of FOXO1 target genes *Murf1* and *Mafbx* (**D-E**). There was no significant effect of FGF21 treatment on glucocorticoid receptor (*Nr3c1*) expression (**F**). Tukey’s HSD posthoc test, **\***<0.05, **\*\***<0.01; N= 9-13/ group.

### FGF21 treatment decreases muscle protein synthesis

To directly measure the effect of FGF21 on muscle protein synthesis we performed stable isotope labeling using deuterium oxide (D2O) (60). Female mice were lighter than male littermates [P (sex) < 0.001] **(Fig. 6A)**. Accordingly, female mice had significantly smaller muscles [P (sex) < 0.001] **(Fig. 6B-C)**. FGF21 treated mice tended to have decreased TA mass [P (treatment) < 0.1] and this was significant when data from mice euthanized after only 1 day of treatment were removed from the model [P (treatment) < 0.01] **(Fig. 6C).** The rate of protein synthesis was greater in female mice compared to males [P (sex) < 0.001]. The effect of FGF21 treatment depended on sex [P (treatment x sex) < 0.05], such that FGF21 significantly decreased the rate of muscle protein synthesis in female TA [Tukey’s posthoc, p< 0.05] **(Fig. 6D)**. These findings support that pharmacologic treatment with FGF21 reduces muscle protein synthesis, and that female mice are more sensitive to this outcome compared to male littermates.

**Figure 6.**
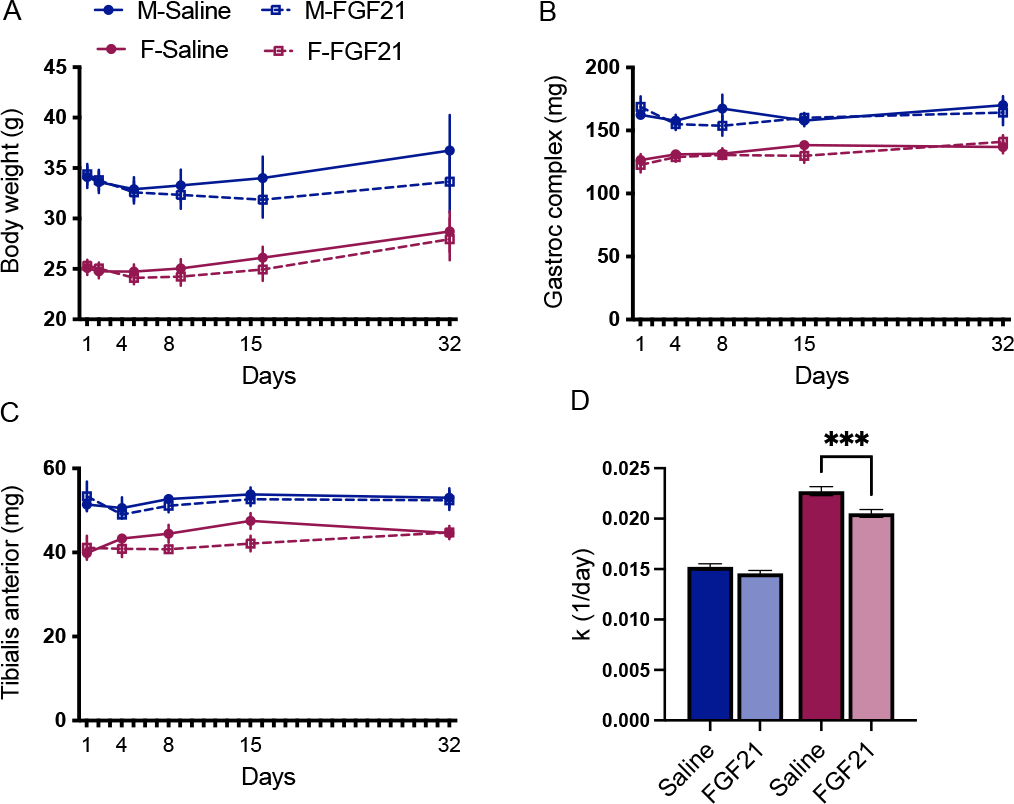
*FGF21 decreased muscle protein synthesis in female mice.* Mice were treated with FGF21 (0.2 mg/kg/d) or saline for 1, 4, 15, or 32 days, while we performed stable isotope labeling with deuterium oxide to measure protein turnover. Female mice (maroon) were lighter than males (blue) [**A**, P (sex) < 0.001]. Female mice had significantly smaller gastrocnemius (**B**) and tibialis anterior (**C**) muscles compared to males [p (sex) < 0.001]. FGF21 treated mice had decreased TA weight [**C**, P (treatment) < 0.01]. The effect of FGF21 treatment on the rate of protein synthesis depended on sex [**D**, P (treatment x sex) < 0.05], such that FGF21 treatment significantly reduced the rate of muscle protein synthesis in female TA, Tukey’s HSD posthoc test, p< 0.05]. n= 3-6 mice per sex/ treatment/ timepoint.

## Discussion

FGF21 is secreted in response to dietary amino acid restriction or imbalance, and other nutritional and metabolic stressors. In turn, FGF21 acts via the nervous system to increase amino acid availability by altering feeding behavior, and shifting macronutrient selection towards increased consumption of dietary protein (8,10,61,62). Here we report that pharmacologic administration of FGF21 also increased systemic amino acids by accessing skeletal muscle, to-gether with activation of the HPA axis and glucocorticoid receptor target genes in skeletal muscle. These findings provide additional insight into the regulation of protein and amino acid homeostasis by FGF21.

In three independent experiments, a short course of treatment with FGF21 either decreased the wet weight of the TA muscle, or reduced TA muscle fiber cross sectional area, in female mice. We used 0.2 mg/kg i.p. or s.c. FGF21 daily. This is a relatively low dose, since most pre-clinical experiments deliver 1 mg/kg/d i.p., but it is nonetheless effective to cause body weight and body fat loss, and to reduce hepatic steatosis in male mice, and it improves glucose tolerance in both sexes (38,63). Because of these beneficial effects on lipid and carbohydrate metabolism, several FGF21 analogues have been developed for human use in treating cardiometabolic diseases (23,35–37)—highlighting the potential translational relevance of our findings. FGF21 analogues are clinically effective to improve fatty liver, though they have performed poorly for body fat loss and glucose control in humans. Changes in muscle mass have not typically been reported. However, in at least one recent clinical trial, the FGF21 analogue Efruxifermin appears to cause loss of lean body mass, since it significantly decreases body weight by an average 3.3 kg while significantly increasing body fat by an average 2.3% (64). FGF21 significantly decreased total lean mass in several animal models but this was not been explored further (34,65–69). Conversely, in another study, when a very low dose of FGF21 (0.1 mg/kg) was administered to male mice for 7 days, no effect on skeletal muscle mass or fiber CSA was observed (70). Considering the present findings, future clinical trials should explicitly consider the effect of FGF21 analogues on muscle mass, with the explicit inclusion of sex as a biological variable potentially influencing this outcome.

To begin understanding the potential physiologic and molecular mechanisms by which pharmacologic FGF21 promotes muscle atrophy, we measured activation of the HPA axis. FGF21 can cross the blood-brain barrier and is present in both human and rodent cerebrospinal fluid (71,72). Moreover, FGFR1 is expressed in CRH neurons of the PVN of the hypothalamus (32). Previous studies by us and others show FGF21 activates the HPA axis (31–33). Intra-PVN injection of recombinant FGF21 increases *Crh* mRNA, plasma adrenocorticotropic hormone (ACTH), and corticosterone (32). Accordingly, the FGF21-treated mice in this study had greater hypothalamic *Crh* mRNA expression, despite no differences in either *Fgfr1* or *Klb*, and circulating corticosterone was significantly increased. FGF21 treatment induced adrenal hypertrophy, indicating chronic stimulation of the adrenal cortex by ACTH. Chronic activation of the HPA axis, or chronic treatment with synthetic glucocorticoids, causes skeletal muscle atrophy and weakness via muscle glucocorticoid receptor (GR) signaling (22). Type 2B “fast” muscle fibers are particularly sensitive to glucocorticoids, due to their higher expression of GR. The TA muscle has a high percentage of Type 2B fibers, and is more sensitive to glucocorticoid-induced atrophy (13), perhaps contributing to the more robust effect of FGF21 on TA vs the gastrocnem-ius-soleus complex.

We found FGF21-induced HPA activation was mirrored by increased expression of *FoxO1, Redd1*, and *Klf15.* These genes are direct and downstream targets of the GR in TA and are thought to decrease protein synthesis by inhibiting mTOR (22,73). When we used stable isotope labeling to directly measure protein synthesis across 32 days of FGF21 treatment, we found FGF21 treated female mice had a decreased rate of muscle protein synthesis. Together these outcomes are consistent with a non-cell autonomous mechanism by which FGF21 induces muscle atrophy, but loss-of-function experiments will be necessary to explicitly determine the role of glucocorticoid receptor.

In these experiments, the effects of pharmacologic FGF21 to reduce mass of the tibialis anterior muscle, decrease protein synthesis, activate the HPA axis, and increase plasma amino acids, were preferentially observed among female mice compared to male littermates. In a recent study, we found the effects of pharmacologic FGF21 to increase lipid catabolism were also sex-dependent, but regarding lipid catabolism, *male* mice were more affected compared to female littermates (38). When we analyzed metabolic pathways associated with FGF21 treatment, we found the five most differently enriched KEGG pathways in females were associated with amino acid metabolism while the five most differently enriched KEGG pathways in males were associated with lipid metabolism. Taken together, these outcomes are consistent with the idea that protein sparing during metabolic stress depends on lipid fuel availability (74,75). That is, greater sensitivity to FGF21-induced lipid catabolism in males may be protein sparing.

Because FGF21 acts in the brain to cause body weight and body fat loss and to increase plasma glucocorticoids in (male) rodents (31–33,76), we reasoned that i.c.v. administration of FGF21 would recapitulate its systemic effects. We used a very low dose that was previously demonstrated to increase HPA axis activity, without efflux to the periphery (32). After a short course of continuous i.c.v. administration, FGF21-treated mice lost more weight than saline-treated controls, and this was driven primarily by its effect in males. Likewise, FGF21-treated male mice, but not female littermates, had smaller inguinal white fat pads than controls. In this experiment there was no significant effect of FGF21 treatment on muscle mass, possibly reflecting the smaller sample size. However histological analysis of fiber CSA showed that FGF21-treated mice had smaller muscle fibers, which was driven primarily by its effect in females. The findings mirror effects observed with systemic treatment, again raising the possibility that greater sensitivity to FGF21-induced lipid catabolism in males may be protein sparing.

Our findings agree with a long literature demonstrating the HPA axis response to stressors are more pronounced in females (41,42), and reports that the effect of glucocorticoids on muscle atrophy are more pronounced in skeletal muscle of females compared to males (43,77). This sex-difference likely results from sex differences in circulating androgens (78,79). For example, early work from Mayer and Rosen showed competitive inhibition of the GR by androgens contributes to their anabolic effect in skeletal muscle (80). Thus, castration increased the binding of labeled dexamethasone to GR in male rats. When both castration and adrenalectomy were performed, there was an inverse relationship between the amount of labeled dexamethasone bound to the GR and the amount of testosterone administered to the animal (81). Future studies will be needed determine the relative contributions of sex hormones and/or chromosomes to these differential outcomes.

The current work focused only on effects of pharmacologic FGF21; when and how endogenously produced FGF21 may affect skeletal muscle mass is also an important topic that requires further investigation. FGF21 was initially identified as a hepatokine, but it can be secreted by other metabolic organs and tissues, including skeletal muscle, in response to intracellular stress (82–85). For example, in a recent study, *Fgf21* mRNA from skeletal muscle was increased roughly 4-fold during a prolonged fast (86) and skeletal muscle specific FGF21-null mice were partly protected against fasting-induced fat loss and muscle atrophy— with parallel differences in muscle protein synthesis and autophagy pathways. Sex of the animals was not reported in that study (86). Likewise, plasma FGF21 is increased, and skeletal muscle mass is decreased in mice lacking the mitochondrial protein Optic Atrophy 1 (OPA1), and these outcomes were diminished among mice lacking both *Fgf21* and *Opa1* in muscle (87). That study used both males and females, but the sex of animals in any given experiment was not reported. Thus, endogenous skeletal muscle FGF21 contributes to muscle atrophy in some conditions. It is not known if this represents a paracrine or endocrine action, since the first order KLB: FGFR1-expressing cells mediating these outcomes were not yet identified. In humans, plasma FGF21 is increased among individuals with diabetes, fatty liver disease, sarcopenia, and mitochondrial myopathies. These physiologic conditions are all associated with muscle atrophy (88–92), but it is not yet known if the increased FGF21 represents a mechanism for muscle loss in these conditions or is simply a biomarker. These are important topics for future research.

## Conclusion

Here we find pharmacologic FGF21 decreases muscle protein synthesis, induces skeletal muscle atrophy, and increases plasma amino acids in female mice. FGF21 is secreted downstream of the intracellular stress response pathway and is thought to act as a neuroendocrine signal of dietary protein and/or amino acid restriction (26). In turn, we (8,62) and others (10) have recently shown that FGF21 can alter feeding behavior and macronutrient selection to increase the consumption of protein, restoring systemic homeostasis. The present studies identify an additional mechanism by which FGF21 can increase systemic amino acid availability— by accessing skeletal muscle. These findings highlight a potential physiologic role for FGF21 in systemic protein metabolism, and they may inform the proposed clinical use of FGF21 analogues for the treatment of cardiometabolic disease.

## ACKNOWLEDGEMENTS

This work was supported by the National Institutes of Health R01DK121035 to KKR, and F31DK124080 to ATC. KRL was supported by a University of California (UC) Davis Floyd and Mary Schwall Dissertation Fellowship. ATC was supported by the National Center for Advancing Translational Sciences, NIH, through grant number UL1 TR001860 and linked award TL1 TR001861. KASS was supported by the National Institute of General Medical Sciences of the NIH T32GM007377. DJ was supported by the National Institute of Arthritis and Musculoskeletal and Skin Diseases funded training program in Musculoskeletal Health Research T32AR079099. Sequencing was carried out at the UC Davis Genome Center DNA Technologies and Expression Analysis Core, supported by NIH Shared Instrumentation Grant 1S10OD010786-01. We thank Fredrick F. Peelor (OMRF) and Yi-Je Chen (UC Davis) for excellent technical assistance.

## CITATIONS

1. Bröer S, Bröer A. Amino acid homeostasis and signalling in mammalian cells and organisms. Biochemical Journal 2017. doi:10.1042/BCJ20160822.

2. Felig P, Owen OE, Wahren J, Cahill GF. Amino acid metabolism during prolonged starvation. The Journal of clinical investigation 1969;48(3):584–594.

3. Kalhan SC, Uppal SO, Moorman JL, Bennett C, Gruca LL, Parimi PS, Dasarathy S, Serre D, Hanson RW. Metabolic and genomic response to dietary isocaloric protein restriction in the rat. J Biol Chem 2011;286(7):5266–5277.

4. **DiBattista D, Holder MD**. Enhanced preference for a protein-containing diet in response to dietary protein restriction. Appetite 1998;30(3):237–254.

5. **Hill CM, Qualls-Creekmore E, Berthoud HR, Soto P, Yu S, McDougal DH, Münzberg H, Morrison CD**. FGF21 and the Physiological Regulation of Macronutrient Preference. Endocrinology 2020;161(3). doi:10.1210/endocr/bqaa019.

6. **Larson KR, Chaffin ATB, Goodson ML, Fang Y, Ryan KK**. Fibroblast Growth Factor-21 Controls Dietary Protein Intake in Male Mice. Endocrinology 2019;160(5):1069–1080.

7. **Morrison CD, Reed SD, Henagan TM**. Homeostatic regulation of protein intake: In search of a mechanism. American Journal of Physiology - Regulatory Integrative and Comparative Physiology 2012. doi:10.1152/ajpregu.00609.2011.

8. **Wu C-T, Larson KR, Goodson ML, Ryan KK**. Fibroblast growth factor 21 and dietary macronu-trient intake in female mice. Physiol Behav 2022;257:113995.

9. **Solon-Biet SM, Cogger VC, Pulpitel T, Heblinski M, Wahl D, McMahon AC, Warren A, Dur-rant-Whyte J, Walters KA, Krycer JR, Ponton F, Gokarn R, Wali JA, Ruohonen K, Co-nigrave AD, James DE, Raubenheimer D, Morrison CD, Le Couteur DG, Simpson SJ**. Defining the Nutritional and Metabolic Context of FGF21 Using the Geometric Framework. Cell Metab 2016;24(4):555–565.

10. **Hill CM, Laeger T, Dehner M, Albarado DC, Clarke B, Wanders D, Burke SJ, Collier JJ, Qualls-Creekmore E, Solon-Biet SM, Simpson SJ, Berthoud H-R, Münzberg H, Morrison CD**. FGF21 Signals Protein Status to the Brain and Adaptively Regulates Food Choice and Metabolism. Cell Rep 2019;27(10):2934–2947.e3.

11. **Kamata S, Yamamoto J, Kamijo K, Ochiai T, Morita T, Yoshitomi Y, Hagiya Y, Kubota M, Ohkubo R, Kawaguchi M, Himi T, Kasahara T, Ishii I**. Dietary deprivation of each essential amino acid induces differential systemic adaptive responses in mice. Molecular Nutrition and Food Research 2014;58(6):1309–1321.

12. **Bodine SC, Furlow JD**. Glucocorticoids and skeletal muscle. Advances in Experimental Medicine and Biology 2015;872:145–176.

13. **Braun TP, Marks DL**. The regulation of muscle mass by endogenous glucocorticoids. Frontiers in Physiology 2015;6(FEB). doi:10.3389/fphys.2015.00012.

14. **Bodine SC, Latres E, Baumhueter S, Lai VKM, Nunez L, Clarke BA, Poueymirou WT, Panaro FJ, Erqian Na, Dharmarajan K, Pan ZQ, Valenzuela DM, Dechiara TM, Stitt TN, Yan-copoulos GD, Glass DJ**. Identification of ubiquitin ligases required for skeletal Muscle Atrophy. Science 2001;294(5547):1704–1708.

15. **Sandri M, Sandri C, Gilbert A, Skurk C, Calabria E, Picard A, Walsh K, Schiaffino S, Lecker SH, Goldberg AL**. Foxo transcription factors induce the atrophy-related ubiquitin ligase atrogin-1 and cause skeletal muscle atrophy. Cell 2004;117(3):399–412.

16. **Shimizu N, Yoshikawa N, Ito N, Maruyama T, Suzuki Y, Takeda SI, Nakae J, Tagata Y, Nishitani S, Takehana K, Sano M, Fukuda K, Suematsu M, Morimoto C, Tanaka H**. Cross-talk between glucocorticoid receptor and nutritional sensor mTOR in skeletal muscle. Cell Metabolism 2011;13(2):170–182.

17. **Waddell DS, Baehr LM, van den Brandt J, Johnsen S a, Reichardt HM, Furlow JD, Bodine SC**. The glucocorticoid receptor and FOXO1 synergistically activate the skeletal muscle atrophy-associated MuRF1 gene. American journal of physiology. Endocrinology and metabolism 2008;295:E785–E797.

18. **Watson ML, Baehr LM, Reichardt HM, Tuckermann JP, Bodine SC, Furlow JD**. A cell-autonomous role for the glucocorticoid receptor in skeletal muscle atrophy induced by systemic glucocorticoid exposure. AJP: Endocrinology and Metabolism 2012;302(10):E1210– E1220.

19. **Wang H, Kubica N, Ellisen LW, Jefferson LS, Kimball SR**. Dexamethasone represses signaling through the mammalian target of rapamycin in muscle cells by enhancing expression of REDD1. Journal of Biological Chemistry 2006;281(51):39128–39134.

20. **Kaplan SA, Shimizu CS**. Effects of cortisol on amino acids in skeletal muscle and plasma. Endocrinology 1963;72:267–272.

21. **Tipton KD, Hamilton DL, Gallagher IJ**. Assessing the Role of Muscle Protein Breakdown in Response to Nutrition and Exercise in Humans. Sports Medicine 2018. doi:10.1007/s40279-017-0845-5.

22. **Kuo T, Harris CA, Wang J-C**. Metabolic functions of glucocorticoid receptor in skeletal muscle. Molecular and Cellular Endocrinology 2013;380(1):79–88.

23. **Fisher FM, Maratos-Flier E**. Understanding the Physiology of FGF21. Annu Rev Physiol 2016;78:223–241.

24. **Nishimura T, Nakatake Y, Konishi M, Itoh N**. Identification of a novel FGF, FGF-21, preferentially expressed in the liver11The nucleotide sequence data reported in this paper will appear in the DDBJ, EMBL and GenBank nucleotide sequence databases with accession numbers AB021975 and AB025718. Biochimica et Biophysica Acta (BBA) - Gene Structure and Expression 2000;1492(1):203–206.

25. **Badman MK, Pissios P, Kennedy AR, Koukos G, Flier JS, Maratos-Flier E**. Hepatic fibroblast growth factor 21 is regulated by PPARalpha and is a key mediator of hepatic lipid metabolism in ketotic states. Cell Metab 2007;5(6):426–437.

26. **Laeger T, Henagan TM, Albarado DC, Redman LM, Bray GA, Noland RC, Münzberg H, Hutson SM, Gettys TW, Schwartz MW, Morrison CD**. FGF21 is an endocrine signal of protein restriction. J Clin Invest 2014;124(9):3913–3922.

27. **Ogawa Y, Kurosu H, Yamamoto M, Nandi A, Rosenblatt KP, Goetz R, Eliseenkova AV, Mohammadi M, Kuro-o M**. βKlotho is required for metabolic activity of fibroblast growth factor 21. Proc Natl Acad Sci U S A 2007;104(18):7432–7437.

28. **Fon Tacer K, Bookout AL, Ding X, Kurosu H, John GB, Wang L, Goetz R, Mohammadi M, Kuro-o M, Mangelsdorf DJ, Kliewer SA**. Research resource: Comprehensive expression atlas of the fibroblast growth factor system in adult mouse. Mol Endocrinol 2010;24(10):2050–2064.

29. **Potthoff MJ, Inagaki T, Satapati S, Ding X, He T, Goetz R, Mohammadi M, Finck BN, Mangelsdorf DJ, Kliewer SA, Burgess SC**. FGF21 induces PGC-1alpha and regulates carbohydrate and fatty acid metabolism during the adaptive starvation response. Proc Natl Acad Sci U S A 2009;106(26):10853–10858.

30. **Inagaki T, Dutchak P, Zhao G, Ding X, Gautron L, Parameswara V, Li Y, Goetz R, Mohammadi M, Esser V, Elmquist JK, Gerard RD, Burgess SC, Hammer RE, Mangelsdorf DJ, Kliewer SA**. Endocrine Regulation of the Fasting Response by PPARα-Mediated Induction of Fibroblast Growth Factor 21. Cell Metabolism 2007;5(6):415–425.

31. **Owen BM, Ding X, Morgan DA, Coate KC, Bookout AL, Rahmouni K, Kliewer SA, Mangelsdorf DJ**. FGF21 Acts Centrally to Induce Sympathetic Nerve Activity, Energy Expendi-ture, and Weight Loss. Cell Metabolism 2014;20(4):670–677.

32. **Liang Q, Zhong L, Zhang J, Wang Y, Bornstein SR, Triggle CR, Ding H, Lam KSL, Xu A**. FGF21 Maintains Glucose Homeostasis by Mediating the Cross Talk Between Liver and Brain During Prolonged Fasting. Diabetes 2014;63(12):4064–4075.

33. **Ryan KK, Packard AEB, Larson KR, Stout J, Fourman SM, Thompson AMK, Ludwick K, Habegger KM, Stemmer K, Itoh N, Perez-Tilve D, Tschöp MH, Seeley RJ, Ulrich-Lai YM**. Dietary manipulations that induce ketosis activate the HPA axis in male rats and mice: A potential role for fibroblast growth factor-21. Endocrinology 2018;159(1):400–413.

34. **Xu J, Lloyd DJ, Hale C, Stanislaus S, Chen M, Sivits G, Vonderfecht S, Hecht R, Li Y-S, Lind-berg RA, Chen J-L, Jung DY, Zhang Z, Ko H-J, Kim JK, Véniant MM**. Fibroblast growth factor 21 reverses hepatic steatosis, increases energy expenditure, and improves insulin sensitivity in diet-induced obese mice. Diabetes 2009;58(1):250–259.

35. **Sanyal A, Charles ED, Neuschwander-Tetri BA, Loomba R, Harrison SA, Abdelmalek MF, Lawitz EJ, Halegoua-DeMarzio D, Kundu S, Noviello S, Luo Y, Christian R**. Pegbelfermin (BMS-986036), a PEGylated fibroblast growth factor 21 analogue, in patients with non-al-coholic steatohepatitis: a randomised, double-blind, placebo-controlled, phase 2a trial. Lancet 2019;392(10165):2705–2717.

36. **Bao L, Yin J, Gao W, Wang Q, Yao W, Gao X**. A long-acting FGF21 alleviates hepatic steatosis and inflammation in a mouse model of non-alcoholic steatohepatitis partly through an FGF21-adiponectin-IL17A pathway. Br J Pharmacol 2018;175(16):3379–3393.

37. **Lee JH, Kang YE, Chang JY, Park KC, Kim H-W, Kim JT, Kim HJ, Yi H-S, Shong M, Chung HK, Kim KS**. An engineered FGF21 variant, L Y2405319, can prevent non-alcoholic steatohepatitis by enhancing hepatic mitochondrial function. Am J Transl Res 2016;8(11):4750–4763.

38. **Chaffin AT, Larson KR, Huang K-P, Wu C-T, Godoroja N, Fang Y, Jayakrishnan D, Sauza KAS, Sims LC, Mohajerani N, Goodson ML, Ryan KK**. FGF21 controls hepatic lipid metabolism via sex-dependent interorgan crosstalk. JCI Insight 2022;7(19). doi:10.1172/jci.in-sight.155848.

39. **Makarova E, Kazantseva A, Dubinina A, Denisova E, Jakovleva T, Balybina N, Bgatova N, Baranov K, Bazhan N**. Fibroblast Growth Factor 21 (FGF21) Administration Sex-Specifically Affects Blood Insulin Levels and Liver Steatosis in Obese Ay Mice. Cells 2021;10(12):3440.

40. **Rincón-Cortés M, Herman JP, Lupien S, Maguire J, Shansky RM**. Stress: Influence of sex, reproductive status and gender. Neurobiol Stress 2019;10:100155.

41. **Kitay JI**. Sex differences in adrenal cortical secretion in the rat. Endocrinology 1961;68:818–824.

42. **Goel N, Workman JL, Lee TT, Innala L, Viau V**. Sex differences in the HPA axis. Compr Physiol 2014;4(3):1121–1155.

43. **Baehr L, Tunzi M, Bodine S**. Muscle hypertrophy is associated with increases in proteasome activity that is independent of MuRF1 and MAFbx expression. Frontiers in Physiology 2014;5.

44. **Sarruf DA, Thaler JP, Morton GJ, German J, Fischer JD, Ogimoto K, Schwartz MW**. Fibroblast growth factor 21 action in the brain increases energy expenditure and insulin sensitivity in obese rats. Diabetes 2010;59(7):1817–1824.

45. **Reeves PG, Nielsen FH, Fahey GC**. AIN-93 Purified Diets for Laboratory Rodents: Final Report of the American Institute of Nutrition Ad Hoc Writing Committee on the Reformulation of the AIN-76A Rodent Diet. The Journal of Nutrition 1993;123(11):1939–1951.

46. **Miller BF, Robinson MM, Reuland DJ, Drake JC, Peelor FF, Bruss MD, Hellerstein MK, Hamilton KL**. Calorie restriction does not increase short-term or long-term protein synthesis. The journals of gerontology. Series A, Biological sciences and medical sciences 2013;68(5):530–8.

47. **Miller BF, Reid JJ, Price JC, Lin H-JL, Atherton PJ, Smith K**. CORP: The use of deuterated water for the measurement of protein synthesis. J Appl Physiol (1985) 2020;128(5):1163– 1176.

48. **Abbott CB, Lawrence MM, Kobak KA, Lopes EBP, Peelor FF, Donald EJ, Van Remmen H, Griffin TM, Miller BF**. A Novel Stable Isotope Approach Demonstrates Surprising Degree of Age-Related Decline in Skeletal Muscle Collagen Proteostasis. Function (Oxf) 2021;2(4):zqab028.

49. **Drake JC, Bruns DR, Peelor FF, Biela LM, Miller RA, Miller BF, Hamilton KL**. Long-lived Snell dwarf mice display increased proteostatic mechanisms that are not dependent on decreased mTORC1 activity. Aging Cell 2015;14(3):474–482.

50. **Lawrence MM, Van Pelt DW, Confides AL, Hunt ER, Hettinger ZR, Laurin JL, Reid JJ, Peelor FF, Butterfield TA, Dupont-Versteegden EE, Miller BF**. Massage as a mechanotherapy promotes skeletal muscle protein and ribosomal turnover but does not mitigate muscle atrophy during disuse in adult rats. Acta Physiol 2020. doi:10.1111/apha.13460.

51. **Miller BF, Hamilton KL, Majeed ZR, Abshire SM, Confides AL, Hayek AM, Hunt ER, Shipman P, Peelor FF, Butterfield TA, Dupont-Versteegden EE**. Enhanced skeletal muscle regrowth and remodelling in massaged and contralateral non-massaged hindlimb: Anabolic effect of massage on skeletal muscle. J Physiol 2018;596(1):83–103.

52. **Edgar R, Domrachev M, Lash AE**. Gene Expression Omnibus: NCBI gene expression and hybridization array data repository. Nucleic Acids Res 2002;30(1):207–210.

53. **Sud M, Fahy E, Cotter D, Azam K, Vadivelu I, Burant C, Edison A, Fiehn O, Higashi R, Nair KS, Sumner S, Subramaniam S.** Metabolomics Workbench: An international repository for metabolomics data and metadata, metabolite standards, protocols, tutorials and training, and analysis tools. Nucleic Acids Res 2016;44(D1):D463–470.

54. Coskun T, Bina HA, Schneider MA, Dunbar JD, Hu CC, Chen Y, Moller DE, Kharitonenkov A. Fibroblast growth factor 21 corrects obesity in mice. Endocrinology 2008;149(12):6018– 6027.

55. **Ulrich-Lai YM, Figueiredo HF, Ostrander MM, Choi DC, Engeland WC, Herman JP**. Chronic stress induces adrenal hyperplasia and hypertrophy in a subregion-specific manner. American Journal of Physiology-Endocrinology and Metabolism 2006;291(5):E965–E973.

56. **Chaffin AT-B, Fang Y, Larson KR, Mul JD, Ryan KK**. Sex-dependent effects of MC4R genotype on HPA axis tone: implications for stress-associated cardiometabolic disease. Stress 2019:1–10.

57. **Goodson ML, Packard AEB, Buesing DR, Maney M, Myers B, Fang Y, Basford JE, Hui DY, Ulrich-Lai YM, Herman JP, Ryan KK**. Chronic stress and Rosiglitazone increase indices of vascular stiffness in male rats. Physiol. Behav. 2017;172:16–23.

58. **Malendowicz LK**. Sex differences in adrenocortical structure and function. VI. Long-term effect of gonadectomy and testosterone or estradiol replacement on rat adrenal cortex. Endokrinologie 1980;75(3):311–323.

59. **Shimizu N, Yoshikawa N, Ito N, Maruyama T, Suzuki Y, Takeda S, Nakae J, Tagata Y, Nishitani S, Takehana K, Sano M, Fukuda K, Suematsu M, Morimoto C, Tanaka H**. Cross-talk between glucocorticoid receptor and nutritional sensor mTOR in skeletal muscle. Cell Metab 2011;13(2):170–182.

60. **Miller BF, Reid JJ, Price JC, Lin H-JL, Atherton PJ, Smith K**. CORES OF REPRODUCIBILITY IN PHYSIOLOGY: THE USE OF DEUTERATED WATER FOR THE MEASUREMENT OF PROTEIN SYNTHESIS. Journal of Applied Physiology 2020:japplphysiol.00855.2019.

61. **Wu C-T, Chaffin AT, Ryan KK**. Fibroblast Growth Factor 21 Facilitates the Homeostatic Control of Feeding Behavior. J Clin Med 2022;11(3):580.

62. **Larson KR, Chaffin AT-B, Goodson ML, Fang Y, Ryan KK**. Fibroblast Growth Factor-21 Controls Dietary Protein Intake in Male Mice. Endocrinology 2019;160(5):1069–1080.

63. **Hale C, Chen MM, Stanislaus S, Chinookoswong N, Hager T, Wang M, Véniant MM, Xu J**. Lack of overt FGF21 resistance in two mouse models of obesity and insulin resistance. Endocrinology 2012;153(1):69–80.

64. **Harrison SA, Ruane PJ, Freilich BL, Neff G, Patil R, Behling CA, Hu C, Fong E, de Temple B, Tillman EJ, Rolph TP, Cheng A, Yale K**. Efruxifermin in non-alcoholic steatohepatitis: a ran-domized, double-blind, placebo-controlled, phase 2a trial. Nat Med 2021;27(7):1262– 1271.

65. **Laeger T, Baumeier C, Wilhelmi I, Würfel J, Kamitz A, Schürmann A**. FGF21 improves glucose homeostasis in an obese diabetes-prone mouse model independent of body fat changes. Diabetologia 2017;60(11):2274–2284.

66. **Emanuelli B, Vienberg SG, Smyth G, Cheng C, Stanford KI, Arumugam M, Michael MD, Adams AC, Kharitonenkov A, Kahn CR**. Interplay between FGF21 and insulin action in the liver regulates metabolism. J Clin Invest 2014;124(2):515–527.

67. **Thompson KE, Guillot M, Graziano MJ, Mangipudy RS, Chadwick KD**. Pegbelfermin, a PEGylated FGF21 analogue, has pharmacology without bone toxicity after 1-year dosing in skeletally-mature monkeys. Toxicol Appl Pharmacol 2021;428:115673.

68. **Stanislaus S, Hecht R, Yie J, Hager T, Hall M, Spahr C, Wang W, Weiszmann J, Li Y, Deng L, Winters D, Smith S, Zhou L, Li Y, Véniant MM, Xu J**. A Novel Fc-FGF21 With Improved Resistance to Proteolysis, Increased Affinity Toward β-Klotho, and Enhanced Efficacy in Mice and Cynomolgus Monkeys. Endocrinology 2017;158(5):1314–1327.

69. **Liu C, Schönke M, Zhou E, Li Z, Kooijman S, Boon MR, Larsson M, Wallenius K, Dekker N, Barlind L, Peng X-R, Wang Y, Rensen PCN**. Pharmacological treatment with FGF21 strongly improves plasma cholesterol metabolism to reduce atherosclerosis. Cardiovasc Res 2022;118(2):489–502.

70. Benoit B, Meugnier E, Castelli M, Chanon S, Vieille-Marchiset A, Durand C, Bendridi N, Pesenti S, Monternier P-A, Durieux A-C, Freyssenet D, Rieusset J, Lefai E, Vidal H, Ruzzin J. Fibroblast growth factor 19 regulates skeletal muscle mass and ameliorates muscle wasting in mice. Nat Med 2017;23(8):990–996.

71. **Lewis JE, Ebling FJP, Samms RJ, Tsintzas K**. Going Back to the Biology of FGF21: New In-sights. Trends in Endocrinology & Metabolism 2019;30(8):491–504.

72. **Hsuchou H, Pan W, Kastin AJ**. The fasting polypeptide FGF21 can enter brain from blood. Peptides 2007;28(12):2382–2386.

73. **Brugarolas J, Lei K, Hurley RL, Manning BD, Reiling JH, Hafen E, Witters LA, Ellisen LW, Kaelin WG**. Regulation of mTOR function in response to hypoxia by REDD1 and the TSC1/TSC2 tumor suppressor complex. Genes Dev. 2004;18(23):2893–2904.

74. **Lowell BB, Goodman MN**. Protein sparing in skeletal muscle during prolonged starvation. Dependence on lipid fuel availability. Diabetes 1987;36(1):14–19.

75. **Jagoe RT, Engelen MPKJ**. Muscle wasting and changes in muscle protein metabolism in chronic obstructive pulmonary disease. Eur Respir J Suppl 2003;46:52s–63s.

76. **Douris N, Stevanovic DM, Fisher FM, Cisu TI, Chee MJ, Nguyen NL, Zarebidaki E, Adams AC, Kharitonenkov A, Flier JS, Bartness TJ, Maratos-Flier E**. Central Fibroblast Growth Factor 21 Browns White Fat via Sympathetic Action in Male Mice. Endocrinology 2015;156(7):2470–2481.

77. Bodine SC, Furlow JD. Glucocorticoids and Skeletal Muscle. In: Wang J-C, Harris C, eds. Glucocorticoid Signaling: From Molecules to Mice to Man. Advances in Experimental Medicine and Biology. New York, NY: Springer; 2015:145–176.

78. **Brouillette J, Rivard K, Lizotte E, Fiset C**. Sex and strain differences in adult mouse cardiac repolarization: importance of androgens. Cardiovascular Research 2005;65(1):148–157.

79. **Handelsman DJ, Hirschberg AL, Bermon S**. Circulating Testosterone as the Hormonal Basis of Sex Differences in Athletic Performance. Endocrine Reviews 2018;39(5):803–829.

80. **Mayer M, Rosen F**. Interaction of glucocorticoids and androgens with skeletal muscle. Metabolism 1977;26(8):937–962.

81. **Mayer M, Rosen F**. Interaction of anabolic steroids with glucocorticoid receptor sites in rat muscle cytosol. American Journal of Physiology-Legacy Content 1975;229(5):1381–1386.

82. **Keipert S, Ost M, Johann K, Imber F, Jastroch M, van Schothorst EM, Keijer J, Klaus S**. Skeletal muscle mitochondrial uncoupling drives endocrine cross-talk through the induction of FGF21 as a myokine. American Journal of Physiology-Endocrinology and Metabolism 2014;306(5):E469–E482.

83. **Miyake M, Nomura A, Ogura A, Takehana K, Kitahara Y, Takahara K, Tsugawa K, Miyamoto C, Miura N, Sato R, Kurahashi K, Harding HP, Oyadomari M, Ron D, Oyadomari S**. Skeletal muscle-specific eukaryotic translation initiation factor 2α phosphorylation controls amino acid metabolism and fibroblast growth factor 21-mediated non-cell-autonomous energy metabolism. FASEB J 2016;30(2):798–812.

84. **Kim KH, Jeong YT, Oh H, Kim SH, Cho JM, Kim Y-N, Kim SS, Kim DH, Hur KY, Kim HK, Ko T, Han J, Kim HL, Kim J, Back SH, Komatsu M, Chen H, Chan DC, Konishi M, Itoh N, Choi CS, Lee M-S**. Autophagy deficiency leads to protection from obesity and insulin resistance by inducing Fgf21 as a mitokine. Nat Med 2013;19(1):83–92.

85. **Guridi M, Tintignac LA, Lin S, Kupr B, Castets P, Rüegg MA**. Activation of mTORC1 in skeletal muscle regulates whole-body metabolism through FGF21. Sci. Signal. 2015;8(402). doi:10.1126/scisignal.aab3715.

86. **Oost LJ, Kustermann M, Armani A, Blaauw B, Romanello V**. Fibroblast growth factor 21 controls mitophagy and muscle mass. J Cachexia Sarcopenia Muscle 2019;10(3):630–642.

87. **Tezze C, Romanello V, Desbats MA, Fadini GP, Albiero M, Favaro G, Ciciliot S, Soriano ME, Morbidoni V, Cerqua C, Loefler S, Kern H, Franceschi C, Salvioli S, Conte M, Blaauw B, Zampieri S, Salviati L, Scorrano L, Sandri M**. Age-Associated Loss of OPA1 in Muscle Impacts Muscle Mass, Metabolic Homeostasis, Systemic Inflammation, and Epithelial Senescence. Cell Metab 2017;25(6):1374–1389.e6.

88. **Koo BK, Kim D, Joo SK, Kim JH, Chang MS, Kim BG, Lee KL, Kim W**. Sarcopenia is an independent risk factor for non-alcoholic steatohepatitis and significant fibrosis. Journal of Hepatology 2017;66(1):123–131.

89. Mesinovic J, Zengin A, De Courten B, Ebeling PR, Scott D. Sarcopenia and type 2 diabetes mellitus: a bidirectional relationship. Diabetes, Metabolic Syndrome and Obesity: Targets and Therapy 2019;12:1057–1072.

90. Hill C, Tallis J. Is obesity a risk factor for skeletal muscle ageing? Aging (Albany NY) 2019;11(8):2183–2184.

91. Kim G, Kim JH. Impact of Skeletal Muscle Mass on Metabolic Health. Endocrinol Metab 2020;35(1):1–6.

92. Pacifico J, Geerlings MAJ, Reijnierse EM, Phassouliotis C, Lim WK, Maier AB. Prevalence of sarcopenia as a comorbid disease: A systematic review and meta-analysis. Experimental Gerontology 2020;131:110801.

